# Novel genes required for the fitness of *Streptococcus pyogenes* in human saliva

**DOI:** 10.1101/196931

**Authors:** Luchang Zhu, Amelia R. L. Charbonneau, Andrew S. Waller, Randall J. Olsen, Stephen B. Beres, James M. Musser

## Abstract

*Streptococcus pyogenes* (group A *Streptococcus*, or GAS) causes 600 million cases of pharyngitis each year. Despite this considerable disease burden, the molecular mechanisms used by GAS to infect, cause clinical pharyngitis, and persist in the human oropharynx are poorly understood. Saliva is ubiquitous in the human oropharynx and is the first material GAS encounters in the upper respiratory tract. Thus, a fuller understanding of how GAS survives and proliferates in saliva may provide valuable insights into the molecular mechanisms at work in the human oropharynx. We generated a highly saturated transposon insertion mutant library in serotype M1 strain MGAS2221, a strain genetically representative of a pandemic clone that arose in the 1980s and spread globally. The transposon mutant library was exposed to human saliva to screen for GAS genes required for wild-type fitness in this clinically relevant fluid. Using transposon-directed insertion site sequencing (TraDIS), we identified 92 genes required for GAS fitness in saliva. The more prevalent categories represented are genes involved in carbohydrate transport/metabolism, amino acid transport/metabolism, and inorganic ion transport/metabolism. Using six isogenic mutant strains, we confirmed that each of the mutants are significantly impaired for growth or persistence in human saliva *ex vivo*. Mutants with an inactivated *spy0644* (*sptA*) or *spy0646* (*sptC*) gene have especially severe persistence defects. This study is the first use of TraDIS to study bacterial fitness in human saliva. The new information we obtained is valuable for future translational maneuvers designed to prevent or treat human GAS infections.

**IMPORTANCE:** The human bacterial pathogen *Streptococcus pyogenes* (group A streptococcus, GAS) causes more than 600 million cases of pharyngitis annually worldwide, 15 million of which occur in the United States. The human oropharynx is the primary anatomic site for GAS colonization and infection, and saliva is the first material encountered. Using a genome-wide transposon mutant screen, we identified 92 GAS genes required for wild-type fitness in human saliva. Many of the identified genes are involved in carbohydrate transport/metabolism, amino acid transport/metabolism, and inorganic ion transport/metabolism. The new information is potentially valuable for developing novel GAS therapeutics and vaccine research.

## [INTRODUCTION]

Bacterial pathogens have evolved highly specialized molecular strategies to survive and persist in diverse host niches (1, 2). Understanding the molecular mechanisms contributing to bacterial fitness in human environments is valuable for developing therapeutic strategies to treat and potentially prevent infections. *Streptococcus pyogenes* (group A streptococcu*s*, or GAS) is a significant human pathogen that causes extensive health and economic impact globally (3). The human oropharynx is the primary anatomic site for GAS colonization and infection (3-5). This pathogen causes 600 million cases of pharyngitis annually worldwide, 15 million of which occur in the United States (3). The annual direct health care costs associated with GAS pharyngitis are estimated to be 2 billion dollars annually in the United States alone (3, 5). The organism is also responsible for an additional 100 million cases of other human infections each year, many of which occur after initial colonization of the oropharynx (3). These additional infections include acute rheumatic fever and subsequent rheumatic heart disease, and as a consequence is the most common cause of preventable pediatric heart disease globally (3, 6). The majority of cases of rheumatic fever occur following human upper respiratory tract infection. Despite the extensive toll on human health, the molecular mechanisms used by GAS to successfully colonize, cause acute pharyngitis and persist in the human oropharynx remain largely unknown or poorly understood (7, 8). This lack of knowledge constitutes a critical knowledge gap in our understanding of GAS pathogenesis, and thus represents an opportunity for enhanced understanding of the molecular mechanisms at work during the critical initial pathogen interaction with the human host.

The oropharynx is the primary site of entry for GAS into the body and the major portal of person-to-person transmission (9-11). Several observations have stimulated our interest in studying the molecular genetic basis of interaction between human saliva and GAS. Saliva is ubiquitous in the human oropharynx and is the first host material contacted by GAS in its common transmission cycle. Compared to individuals with clinical pharyngitis who lack GAS in their saliva, patients with GAS present in their saliva are more likely to transmit the organism to a new host by aerosolization (10-12). Thus, understanding how GAS survives and proliferates in saliva and the oropharynx may provide valuable insights into the molecular mechanisms underlying successful bacterial interaction in this niche.

Previous studies addressing GAS-saliva interactions have identified several factors that contribute to bacterial fitness (13-18), but knowledge is limited. The studies conducted by Sitkiewicz et al. (13) and Virtaneva et al. (14, 15) investigated gene responses of a serotype M1 GAS strain grown in human saliva *ex vivo* (16). Subsequently, Shelburne et al (17) identified a key two-component transcriptional regulatory system (SptR/S) of previously unknown function that plays central role in optimizing GAS persistence in saliva. The SptR/S two-component system influences multiple GAS metabolic pathways and production of many virulence factors (17). For example, the secreted virulence factors streptococcal inhibitor of complement (Sic) and a potent extracellular cysteine protease (SpeB) made by GAS during growth *ex vivo* in saliva contribute significantly to GAS persistence in this fluid (16). Additional information about GAS interaction in the oropharynx was provided by a 20-monkey study that investigated global transcriptome changes occurring over 86 days of the infection cycle, including initial colonization, acute clinical pharyngitis, and ultimately asymptomatic colonization (15, 18). Taken together, these studies have provided a broad overview of some of the GAS processes at work in the oropharynx; however, much remains to be learned.

Especially lacking is a detailed understanding of the genes required for successful growth and persistence in saliva. To address this important knowledge gap, we conducted a genome-wide screen to identify GAS genes contributing to fitness in human saliva *ex vivo*. Using <u>**tra**</u>nsposon-<u>**d**</u>irected <u>**i**</u>nsertion <u>**s**</u>ite sequencing (TraDIS) (19-28), we generated a highly saturated transposon insertion library (140,249 unique transposon insertions) in serotype M1 reference strain MGAS2221 (29-31). This serotype was used because it is usually the most common cause of pharyngitis and other human infections in western countries (3). Strain MGAS2221 is genetically representative of the pandemic clone that arose in the 1980s and rapidly spread globally (29-32). The transposon mutant library was exposed to human saliva *ex vivo* for 48 hrs to identify GAS genes contributing to fitness over time in this clinically relevant fluid. Saliva studies conducted with six isogenic mutant strains validated the findings. The new information we obtained substantially increases our overall understanding of the molecular genetic basis of pathogen-saliva interactions and is valuable for future translational research designed to treat or prevent human GAS infections.

## RESULTS

### Construction of a highly saturated transposon insertion library in serotype M1 GAS strain MGAS2221

A transposon insertion mutant library was generated using serotype M1 strain MGAS2221 as the parental organism. Strain MGAS2221 was chosen for transposon mutagenesis because (i) it is genetically representative of a pandemic clone that arose in the 1980s and disseminated worldwide (30-32), (ii) MGAS2221 has wild-type alleles of major transcriptional regulators that affect virulence, such as *covR*/*covS*, *ropB*, *mga*, and *rocA*, and (iii) it has been used in many mouse and primate infection studies (31). Using plasmid pGh9:IS*S1* (26), we successfully generated a dense transposon mutant library in strain MGAS2221 containing 140,249 unique transposon insertions. On average, the library contains one transposon insertion every 13 nucleotides. 93.4% of the genes in the MGAS2221 genome (1720 out of 1841) have at least one transposon insertion. The nearly random distribution of transposon insertion and high density of transposon insertions in the *S. pyogenes* genome is illustrated in Fig. 1. By analyzing the mutant library using the tradis_essentiality TraDIS toolkit script (25), we identified 432 genes (∼23.5% of the genes in the genome) that are essential for the serotype M1 GAS strain MGAS2221 in our experimental conditions (40°C, in THY broth supplemented with 0.5 μg/ml erythromycin, see Materials and Methods). The list of identified essential genes is presented in Table S1.

**FIG 1.**
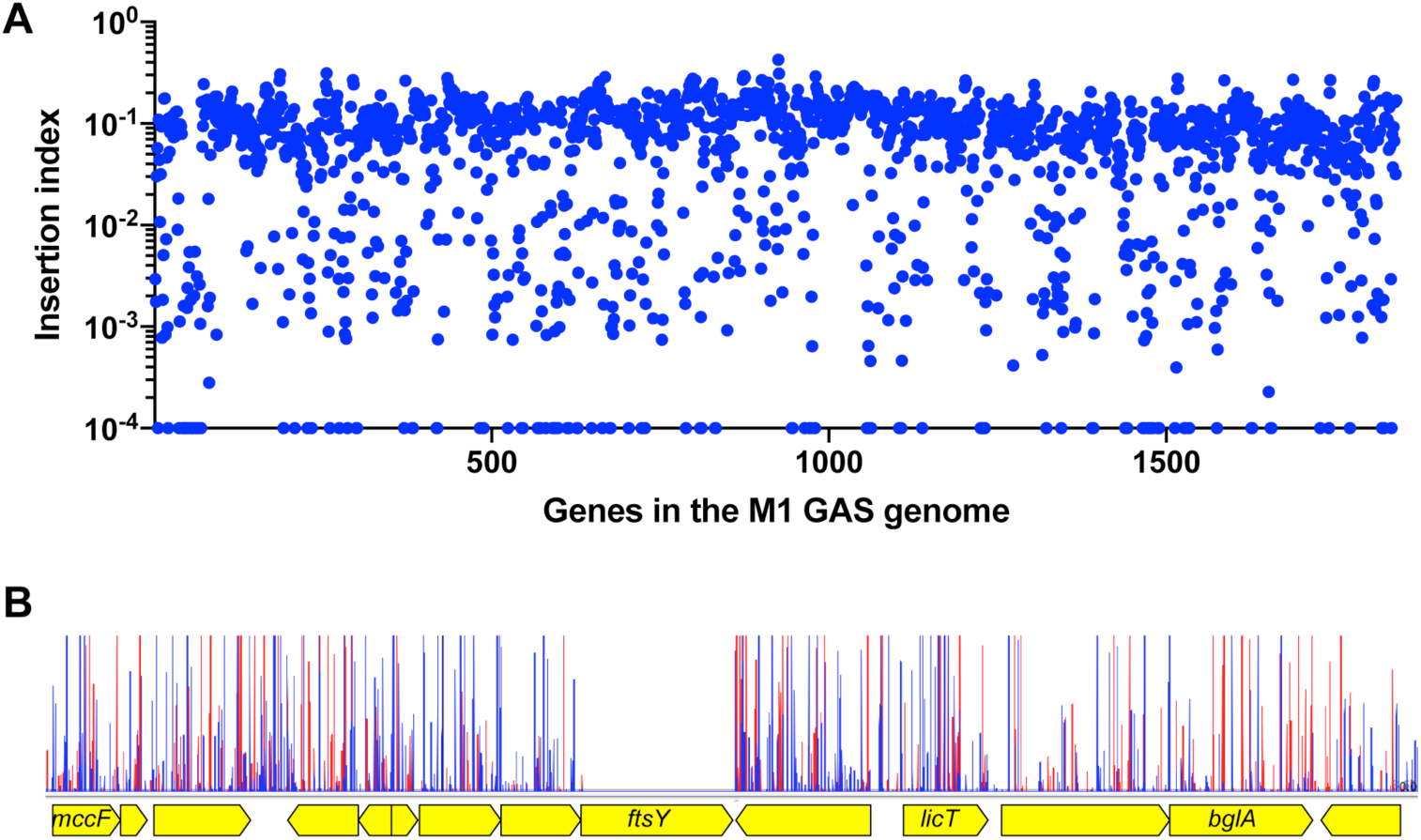
Near-random distribution (A) and high density of transposon insertions (B) of the M1 GAS input mutant library. (A) Insertion index (number of insertion sites divided by gene length) (Y-axis) of each gene (X-axis) in the M1 GAS reference genome. (B) A representative section of the transposon insertion map. As expected, essential gene ftsY has no insertion because it is not represented in the library. Red vertical spikes are forward reads; blue vertical spikes are reverse reads. Read orientation indicates the direction of the transposon insertion.

### Genes contributing to fitness of GAS over time in human saliva

We exposed the transposon mutant library to pooled human saliva *ex vivo* to screen for genes contributing to fitness in this fluid. TraDIS was used to identify genes with significantly altered mutant frequency in the output mutant pools compared to the input pool at 12, 24, and 48 hrs after saliva inoculation. Genes with significantly decreased mutant frequency (fold-change >1.5, and q value < 0.1) in the output mutant pools were regarded as important for saliva growth and persistence, which can be referred to as fitness. To ensure that the statistical power was adequate, genes with fewer than 10 transposon insertions in any of the four input mutant pools were excluded from the analysis, as recommended by van Opijnen et al (33).

We identified 30 (12 hrs), 42 (24 hrs) and 83 (48 hrs) genes with significantly decreased mutant frequencies, providing evidence that these genes contribute to GAS fitness in human saliva (Fig. 2 and Fig. 3, Table S2). In total, 92 genes were identified at the three time points (Fig. 3A, Table S2). Clusters of orthologous groups (COG) classification of the 92 genes showed that numerically, the three more prevalent categories include genes involved in carbohydrate transport and metabolism (*n* = 10 genes), amino acid transport and metabolism (*n* = 8 genes), and inorganic ion transport and metabolism (*n* = 7 genes) (Fig. 3B). Our previously published data from an experimental pharyngitis infection study, involving 20 cynomolgus macaques (15), identified genes expressed during GAS oropharyngeal infection. Of the 92 saliva-fitness genes identified by TraDIS, ∼74% were also expressed during GAS oropharyngeal infection (Table S2). Moreover, many of these 92 genes (e.g., *nagA*, *pstS*, *oppA*, and *malX*) are upregulated during GAS oropharyngeal infection in cynomolgus macaques (15). Together, our results suggest that many of the GAS genes contributing to fitness in saliva *ex vivo* may also contribute to pathogen fitness in the oropharynx of non-human primates (NHPs). However, experiments will be required to directly test this hypothesis.

**FIG 2.**
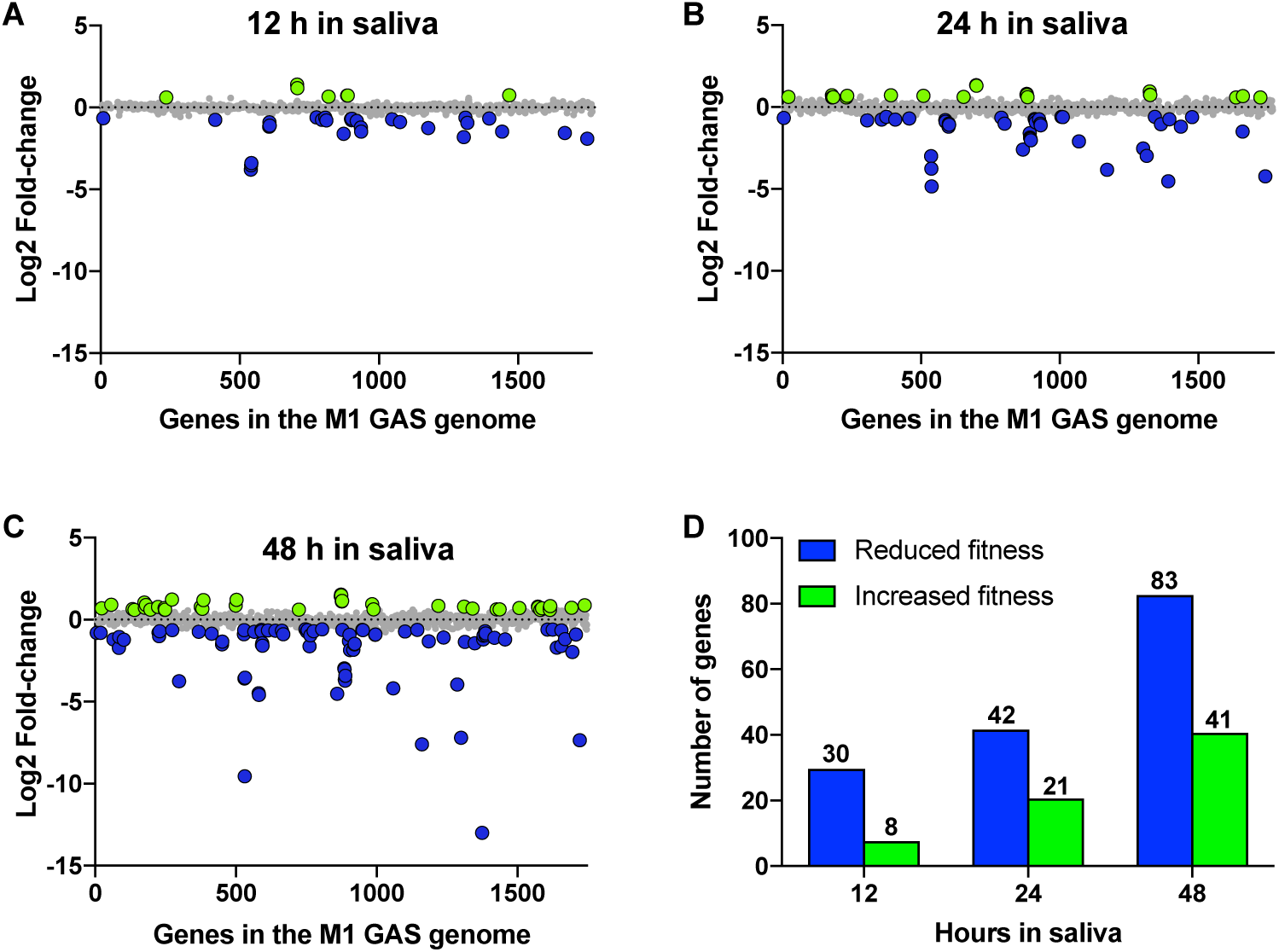
TraDIS analyses of GAS fitness genes during exposure to human saliva *ex vivo*. (A-C) Genome-scale summary of the change in mutant abundance (Y-axis) for each gene (X-axis) in the output mutant pools recovered after 12 hrs, 24 hrs and 48 hrs of incubation in human saliva *ex vivo*. Gene mutations (insertions) conferring significantly decreased (blue circles) or increased (green circles) fitness are highlighted. Insertion mutations that lack a significantly altered fitness phenotype (grey circles) are also indicated. (D) Summary of GAS genes identified to be important for fitness in saliva after indicated period of saliva incubation.

**FIG 3.**
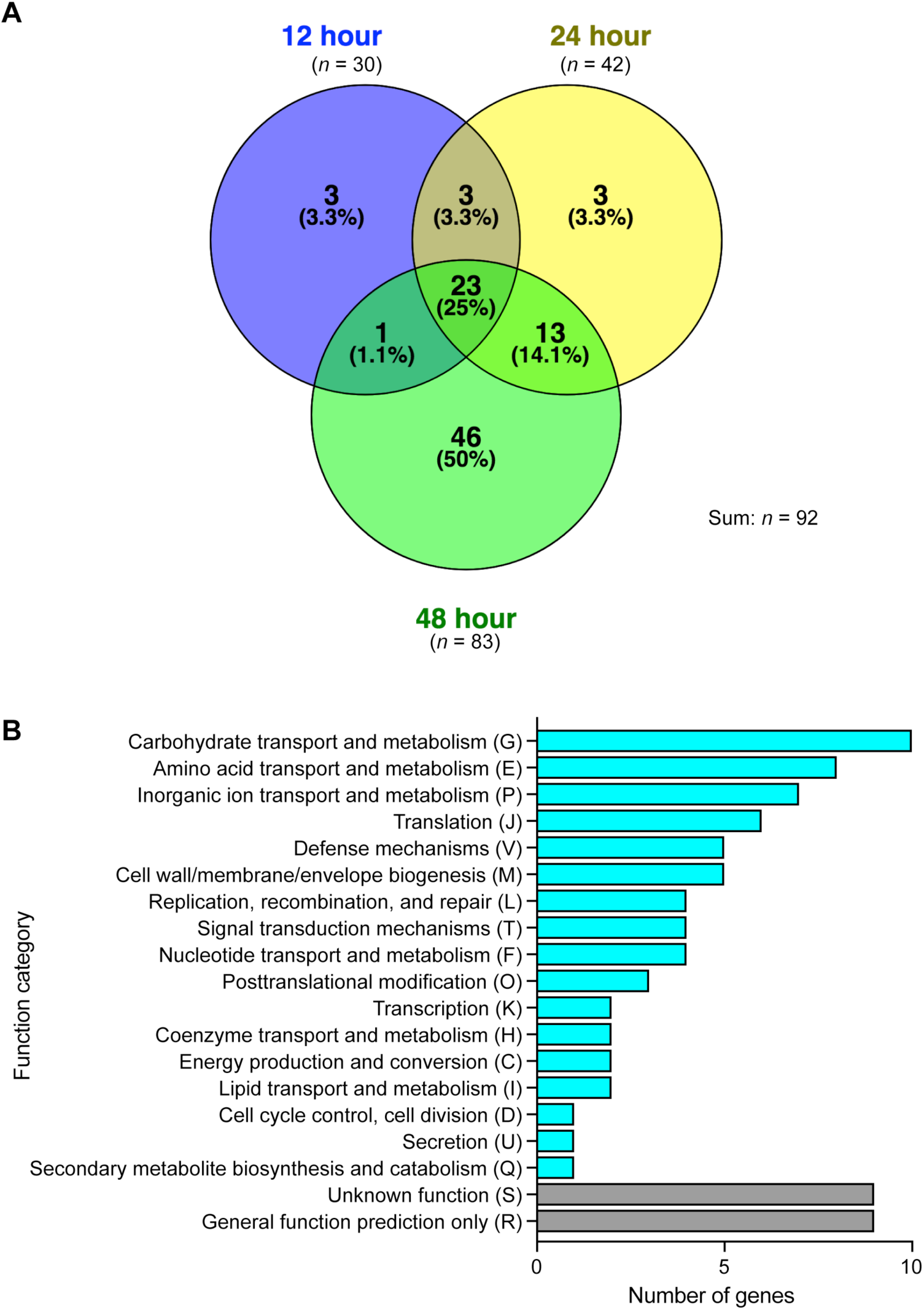
Venn diagram showing number of GAS genes identified to be important for fitness in saliva after indicated period of incubation (A), and functional categorization of the 92 identified GAS saliva-fitness genes (B). Note that in panel A the circles sizes are not proportional to the numbers of genes identified simply to improve visual presentation and clarity.

Additionally, we determined that some genes at 12 hr (n = 8 genes), 24 hr (n = 21 genes) and 48 hr (n = 41 genes) were significantly associated with potentially increased GAS fitness in saliva. The magnitude of the fold-change in these genes was relatively modest compared to the genes that were decreased in fitness (Fig. 2A-C, Table S3).

The fitness score of all genes in the M1 GAS genome (including the genes with less than 10 insertions) with significant change in sequence read counts at the three time points (12 hr, 24 hr and 48 hr) are listed in Table S4, Table S5, and Table S6, respectively.

### Validation of six genes required for wild-type GAS fitness in human saliva *ex vivo*

To validate the TraDIS screen findings, we analyzed the saliva fitness phenotype of each isogenic mutant strain generated from six genes (*spy0644*, s*py0646*, *lacR.1*, *carB*, *nifS1*, and *pstS*) identified in the screen. These six genes were chosen for analysis because (i) they have not been previously shown to participate in GAS fitness in human saliva, (ii) transposon insertions into these genes represented a range of altered fitness fold-change values, (iii) the genes are present in the core genome of all sequenced GAS genomes, (iv) the genes are known to be expressed in the oropharynx of NHPs during experimental infection (15), and (v) these genes participate in a variety of biological pathways: *spy0644* and s*py0646* (putative ABC transporter), *lacR.1* (carbohydrate metabolism), *carB* (pyrimidine and arginine synthesis), *nifS1* (amino acid metabolism) and *pstS* (phosphate transport). To test the hypothesis that inactivating each of these six genes impaired GAS fitness in human saliva *ex vivo*, we used targeted insertional mutagenesis (31, 34, 35) to create isogenic mutant strains from wild-type parental strain MGAS2221. The genome of each isogenic strain was sequenced before use to ensure that no spurious mutations had been introduced during mutant construction. Consistent with our hypothesis, the results (Fig. 4) confirmed that these six isogenic mutant strains had significantly decreased fitness in human saliva compared to parental strain MGAS2221 (Fig. 4). Importantly, we discovered that mutant strains Δ*spy0644* and Δ*spy0646* had severely impaired fitness in saliva (Fig. 4A). Greater than 99% of the Δ*spy0644* and Δ*spy0646* inocula were not present as viable cells at the 24 hr time point post-saliva inoculation (Fig. 4A). Genes *spy0644*, *spy0645*, and *spy0646* likely constitute an operon that encodes an ABC transporter system (Fig. 4A and Table S2). On the basis of genome sequencing of thousands of strains of 20 evolutionarily diverse M protein serotypes commonly causing human infections, these three genes are part of the core genome of GAS (32, 36). That is, these genes are present in all GAS genomes sequenced and moreover, they are highly conserved in genome location and context, and primary amino acid sequence. In addition, homologs of this three-gene region are present in related species of pathogenic streptococci, including *Streptococcus agalactiae, Streptococcus dysgalactiae*, *Streptococcus equi*, *Streptococcus gallolyticus*, *Streptococcus mutans* and others, and are conserved in location downstream of *carB* (Fig. 5). This suggests a functional relationship exists between this ABC transporter and the metabolic activities of CarB. In this regard, we also identified *carB* to be important for wild-type GAS fitness in saliva (Table S2), and confirmed that the Δ*carB* isogenic mutant strain is significantly impaired in persistence in saliva (Fig. 4B). *carB* encodes the large subunit of carbamoylphosphate synthetase (37, 38). Carbamoylphosphate is a precursor for both pyrimidine and arginine synthesis (38). Interestingly, *carB* also was reported to be required for GAS fitness in human blood, a body fluid with a very different chemical composition to saliva (39). These results indicate that pyrimidine and arginine synthesis mediated in part by *carB* contributes to GAS fitness in multiple host niches. The Δ*pstS* mutant strain is also significantly attenuated for growth in human saliva (Fig. 4E). *pstS* encodes a putative phosphate binding protein and is part of a six-gene operon encoding a phosphate uptake system (Fig. 4E). A genome-wide transposon mutagenesis screen found that *pstS* is required for the fitness of *Streptococcus pneumoniae* in human saliva *ex vivo* (40). We also note that *pst* operon genes have been reported to contribute to the virulence of multiple gram-negative pathogens, including *Proteus mirabilis* and *Escherichia coli* (41-45). These results suggest efficient phosphate uptake is important for fitness of multiple human and animal pathogens that must successfully interact with their specific host niches, including saliva in the oropharynx. Growth of the six mutants in rich medium *in vitro* (THY) showed that none of the mutants has severe growth defects (Fig. 4F). To summarize, the saliva growth phenotype of these six isogenic mutant strains strongly reflected the fitness data obtained from the high-throughput TraDIS screen.

**FIG 4.**
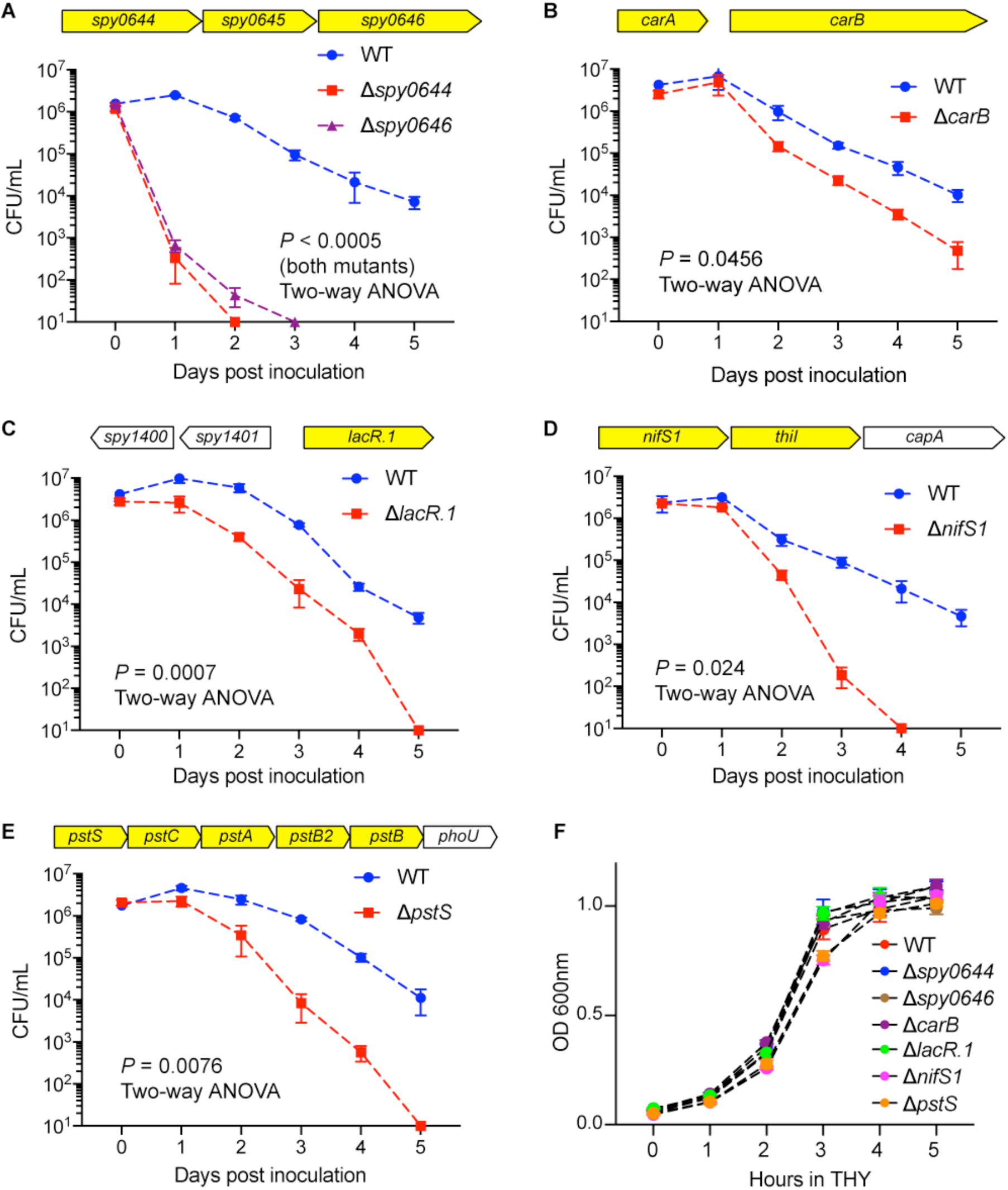
Validation of the findings from the TraDIS saliva screen. The saliva persistence phenotype was determined for each of six GAS isogenic mutant strains (A-E). Highlighted genes (yellow) indicate the putative saliva-fitness genes identified by TraDIS. *P*-values were assessed by repeated-measures 2-way ANOVA. The growth phenotype in rich medium (THY) was also determined for each of six GAS isogenic mutant strains (F).

**FIG 5.**
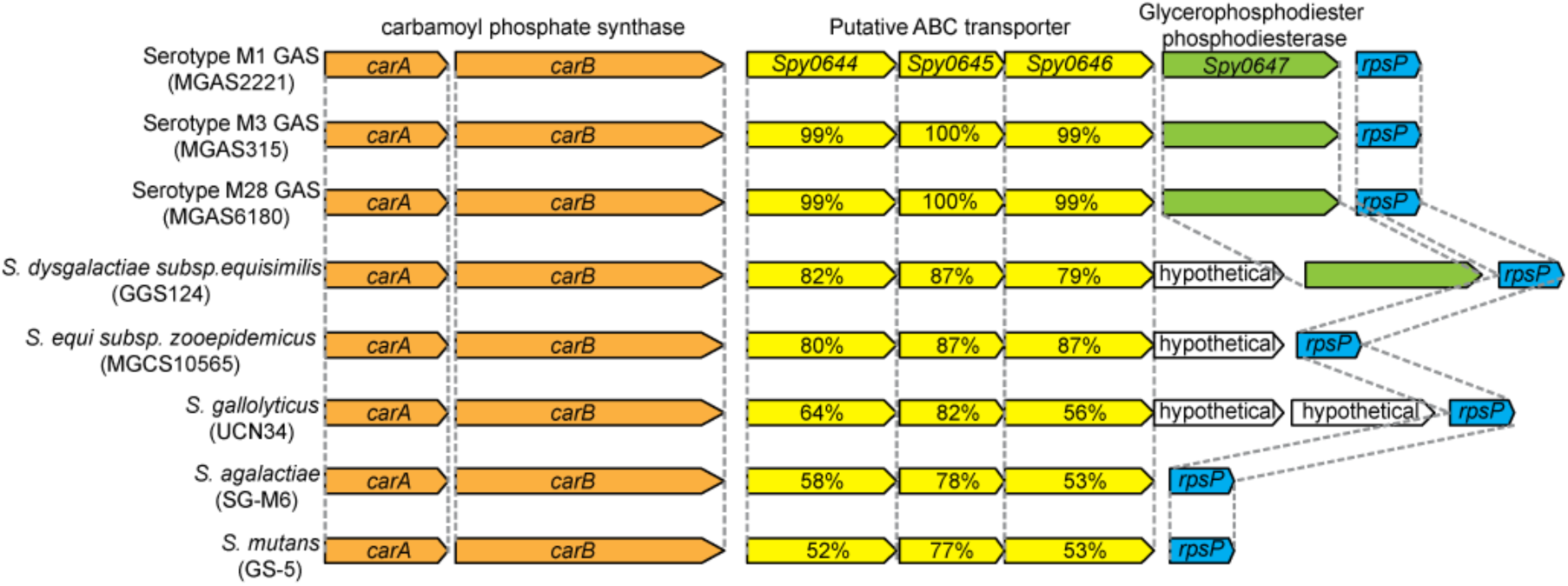
Homologous regions encoding genes *Spy0644*-*Spy0646* in GAS and other bacteria. Yellow arrows represent genes *Spy0644*, *Spy0645*, *Spy0646* and their homologs. Percentages denote amino acid identity compared to serotype M1 GAS strain MGAS2221.

## DISCUSSION

Our results present for the first time a genome-wide view of the GAS genes contributing to pathogen fitness in human saliva, only the second pathogen (40) for which such a screen has been performed. The work also represents the first application of TraDIS to GAS, a human-specific pathogen responsible for greater than 700 million infections each year, including 600 million pharyngitis cases (3).

Some years ago we initiated study of GAS–human saliva molecular interactions (16) with the goal of obtaining new insight into GAS gene activity during the earliest stage of oropharyngeal infection. A common theme that has emerged from many studies (16, 17, 46, 47) is that genes involved in complex carbohydrate catabolism play important roles in growth and persistence in human saliva. Saliva in the human oropharynx contains many nutrients and diverse molecules critical to innate and acquired immunity (48-50). Knowledge gained about how GAS responds to saliva contributes to a broader understanding of host–pathogen interaction and microbial persistence on mucosal surfaces. Expression microarray analysis, immunologic methods, and *in vivo* gene quantification identified a genetic program used by GAS to survive in human saliva *ex vivo* (14, 16, 17). A key discovery was that a two-component regulatory system (TCS) of previously unknown function played a central role in pathogen survival in saliva (17), and revealed an intimate link between metabolism, virulence factor production, and GAS persistence in saliva. However, the strategy used was unable to directly identify the specific genes contributing to GAS persistence in saliva. Our TraDIS analysis discovered that 25 of 92 (27%) genes contributing to fitness in human saliva *ex vivo* are involved in carbohydrate, amino acid and inorganic ion transport and/or metabolism. Inactivation of genes in these categories is likely to significantly impair core metabolic processes such as nutrient acquisition and use. Our results add to the important theme of an intimate linkage between metabolism and GAS persistence in human saliva. We note that a similar spectrum of genes was abundantly represented in an *ex vivo* human saliva transposon mutagenesis screen conducted for *S. pneumoniae* (40). For example, the *opp* operon (oligo peptide transport) and *pst* operon (phosphate uptake) are required for fitness in both GAS and *S. pneumoniae* (40) (Table S2), suggesting certain mechanisms contributing to bacterial fitness in saliva are shared by multiple pathogens in the oropharyngeal niche.

By mining data available from our previously conducted NHP pharyngitis study, we found that a large majority (74%, see Table S2) of the 92 genes discovered herein to contribute to fitness in human saliva also were expressed *in vivo* in the monkey oropharynx (15). Several explanations may account for the lack of evidence for expression of 24% of the genes. First, the NHP study was conducted relatively early in our understanding of the annotation of the genome of the input serotype M1 strain MGAS5005. It is possible that not all of the genome of MGAS5005 was represented on the Affymetrix gene chip used in that study. Second, it is possible that some of the genes were expressed, but at levels too low to detect with the techniques available at that time. Third, it may be that some of the genes in MGAS2221 are expressed at different time points than we used in the two studies, for example farther into the asymptomatic carriage phase (that is, later than day 7), the end point used in our current study. Finally, human saliva *ex vivo* is a different environment than the primate oropharynx, a niche that also contains, for example, host innate and acquired immune cells and epithelial cells. Therefore, it was not unexpected to find potential differences in evidence of gene expression between the two data sets. Despite this, the remarkable 74% gene overlap unambiguously shows that many similar mechanisms are at work on GAS regardless of whether it is exposed to human saliva *ex vivo* or inoculated into the primate oropharynx.

For decades, the analysis of GAS-mediated processes contributing to pharyngitis was restricted predominantly to inferences obtained by evaluating serologic responses to relatively few extracellular molecules that participate in pathogen-host molecular interactions, such as M protein, DNase B, and streptolysin O (51-53). Although important information has been obtained from these descriptive studies, the inability to directly identify large numbers of GAS genes contributing to pharyngitis means that we have a very imprecise understanding of molecular processes contributing to this important human infection. Given the 74% overlap in genes between our TraDIS screen and NHP pharyngitis expression data, it is reasonable to conclude that our study advances understanding of molecular events occurring in the oropharynx, and thereby provides a critical foundation for subsequent molecular pathogenesis studies.

Growth and persistence in human saliva contributes to the ability of GAS to be successfully transmitted by respiratory droplets. The work of Hamburger (9-11) demonstrated that individuals with higher CFUs of GAS in saliva were more likely to transmit the organism to others. This observation implies that any process detrimentally affecting fitness in saliva (such as gene inactivation as done herein) is likely to decrease the probability of successful transmission. It also suggests that mutants with substantial fitness defects are more highly likely to have decreased ability to disseminate and/or cause clinical pharyngitis. The isogenic mutants constructed by inactivation of *spy0644* (herein denoted *sptA*, for streptococcal persistence) and *spy0646* (*sptC*) had the most pronounced growth phenotype of the six isogenic mutant strains during saliva growth *ex vivo*. Of note, the transposon mutants of these two genes had the lowest fitness scores at the early time point (12 hr) of any of the six mutants we tested (Table S2). In contrast, according to the screen, *carB* and *lacR.1* transposon mutants have significantly decreased fitness only at the latest time point (48 hr) (Table S2). Isogenic mutants Δ*carB* and Δ*lacR.1* had a moderate saliva growth phenotype compared to the Δ*sptA* and Δ*sptC* strains (Fig. 4B and C). These findings suggest that a relationship exists between magnitude of the fold-change at an early time point and growth of the resulting isogenic mutant strain in saliva *ex vivo*. However, more data generated with additional isogenic mutant strains is required to rigorously test this idea. If true, use of these data could be an important characteristic used to help triage GAS genes for more in depth analysis, including translation research activities.

Our results show that mutants with insertions in *pst* operon genes are significantly attenuated for growth in human saliva (Fig. 4E, Table S2). The *pst* operon encodes a high-affinity phosphate transporter and has been reported to contribute to bacterial virulence and fitness in a wide variety of pathogens (41, 43-45, 54-56). For example, genome-wide transposon mutagenesis screens found that *pst* operon is required for the fitness of *S. pneumoniae* in human saliva *ex vivo* (40) and mouse lung infections (57). In the oral pathogen *S. mutans*, deleting *pstS* results in decreased production of extracellular polysaccharide, and reduced ability to adhere to saliva-coated surfaces (56). The *pst* operon genes in gram-negative pathogens, have been reported to contribute to the virulence of *Proteus mirabilis* and *Escherichia coli* (41-45, 54). Together, these results suggest efficient phosphate uptake is important for the ability of multiple human and animal pathogens to survive and thrive in their specific host niches, including *S. pyogenes* in human saliva.

The transposon mutagenesis screen study we performed has several limitations. In the human oropharynx, saliva is constantly replenished, whereas in our experimental system a single aliquot of pooled saliva was used for the entire incubation period. A second important limitation is the lack of intact immune cells contributing to innate and acquired immunity in the saliva preparation used. These and other cells (e.g., epithelial) are lacking because the saliva is filter-sterilized prior to use. It is also possible that very small genes were missed in the screen because we excluded genes with fewer than 10 inserts from the analysis, a common practice in transposon mutagenesis studies (33, 58). Overcoming some of these limitations will require experimental infection of an intact animal that faithfully recapitulates all phases of human pharyngitis, such as NHPs. A third limitation is the six isogenic mutants we used to validate the TraDIS results were generated by insertional inactivation. Although the isogenic mutants have no spurious mutations, and their phenotype recapitulated the TraDIS findings, this does not rule out potential polar effects on neighboring genes, especially genes located in the same operon. To overcome this limitation in future follow-up in-depth functional studies on key genes and operons identified in this initial screen, non-polar deletion mutants and complemented mutant strains can be used.

To summarize, by identifying genes contributing to fitness in human saliva *ex vivo*, our work complements important information obtained in other transposon mutant screens that identified GAS genes contributing to fitness in human blood *ex vivo* and mouse soft tissue disease after subcutaneous infection, and for growth *in vitro* (39, 59, 60). It is reasonable to suggest that the information we obtained from this genome-wide screen for genes contributing to fitness in human saliva can be successfully exploited for future pathogenesis investigations.

## MATERIALS AND METHODS

### Bacterial strains and growth conditions

Strain MGAS2221 is genetically representative of a pandemic clone of serotype M1 that arose in the 1980s and has spread worldwide (30-32). Isogenic mutant strains Δ*carB*, Δ*lacR.1*, Δ*sptA*, Δ*sptC*, Δ*nifS1*, Δ*pstS* were derived from parental strain MGAS2221, the organism used for construction of the transposon mutant library. All GAS strains were grown in Todd-Hewitt broth supplemented with 0.2% yeast extract (THY broth) at 37°C with 5% CO_2_.

### Generation of GAS transposon mutant library

A transposon mutant library was generated in strain MGAS2221 using transposon plasmid pGh9:IS*S1* based on a recently described protocol (26). Briefly, pGh9:IS*S1* was transformed into strain MGAS2221 by electroporation (26). A single colony of the transformants was picked and grown overnight at 28°C (permissive temperature) in THY broth supplemented with 0.5 μg/ml erythromycin. The resulting overnight culture was heat shocked at 40°C (nonpermissive temperature) for 3 hrs to permit random transposition and integration of pGh9:IS*S1* into the GAS genome. The GAS cells in the culture were harvested by centrifugation, plated on THY agar supplemented with 0.5 μg/ml erythromycin, and grown overnight at 37°C. The transposon mutant library (i.e., pooled transposon mutants) was collected by washing the colonies off the agar plates with THY broth containing 25% glycerol. The bacterial cell suspension (transposon mutant library) was stored at −80°C. This process was repeated on three separate occasions and the three libraries were pooled.

### Human saliva collection and processing

Human saliva was collected and processed as described previously (16, 17). Briefly, saliva was collected from five healthy donors, pooled, clarified by centrifugation at 45,000 x g for 15 min and sterilized with a 0.20 μm filter (Corning Inc.). The resulting sterile saliva was used for subsequent transposon mutant library screening and individual strain growth.

### Exposure of the transposon mutant library to human saliva *ex vivo*

The transposon mutant library was inoculated into 20 ml of THY and grown at 37°C to mid-exponential phase (OD = 0.5). The bacteria were harvested by centrifugation and washed three times with an equal volume of PBS to remove trace THY broth. 50 μl of the cell suspension in PBS was inoculated into four tubes, each containing 40 ml of filtered saliva (Fig. S1). The four inoculated tubes constitute four biological replicates of the saliva persistence assay (Fig. S1). Immediately after inoculation, 200 μl of the inoculated saliva from each of the four replicate cultures was plated onto four THY plates, and incubated at 37°C for 12 hrs. GAS cells growing on the plates were harvested and represent the composition of the input mutant pools (0 hr mutant pools, *n* = 4). Mutant pools present at 12 hrs, 24 hrs and 48 hrs post-inoculation were recovered by plating 200 μl of the inoculated saliva onto THY agar plates at the aforementioned time points and incubated for 12 hrs at 37°C. The collected mutant pools (4 replicates, 4 time points, *n* = 16) were stored at −80°C for the subsequent TraDIS analysis.

### DNA preparation and massive parallel sequencing

Genomic DNA preparation and DNA sequencing was performed according to procedures described previously for TraDIS analysis, with minor modifications (26). Briefly, genomic DNA of mutant pools collected at the various time points was isolated using DNeasy blood & tissue kit (Qiagen). 2 μg purified genomic DNA was fragmented by incubating with NEBNext dsDNA Fragmentase (New England Biolabs) for 25 min at 37 °C to obtain DNA fragments in the range of 200-1,000 bp. A Y-adaptor (26) (Table S7) was ligated to 1 μg of fragmented DNA using the NEBNext Ultra II DNA library prep kit for Illumina (New England Biolabs). The adaptor-ligated DNA fragments were then purified using AMPure XP beads (Agencourt, Beckman Coulter) and digested with restriction enzyme BamHI for 3 hrs at 37 °C to minimize the mapping of TraDIS reads to the transposon plasmid backbone (26). The resulting DNA was purified, and 100 ng of the purified library DNA was subjected to PCR using the specific IS*S1* primer and one of the 8 indexing PCR primers per DNA library (Table S7), to amplify regions that span the 5' end of IS*S1* and the GAS genomic regions adjacent to the chromosomal location of the transposon. The PCR amplified libraries were sequenced using a single end 76-cycle protocol on a NextSeq550 instrument (Illumina) using a custom Read 1 primer (Table S7) and a custom Index Read sequencing primer (Table S7).

### Processing of DNA sequencing reads and data analysis

The raw Illumina reads obtained from the input and output pools were parsed with FASTX Barcode Splitter (http://hannonlab.cshl.edu/fastx_toolkit/commandline.html). After removing adaptor, low quality reads, and index sequences, PRINSEQ lite version 0.20.4 (http://prinseq.sourceforge.net/) was used to eliminate reads shorter than nucleotides. The resulting sequencing reads were analyzed with the TraDIS toolkit (25) according to previously described methods (26, 61). Briefly, bacteria_tradis was used to trim transposon tag sequence and map the remaining reads to the serotype M1 strain MGAS5005 reference genome. The plot files generated by bacteria_tradis were analyzed by tradis_gene_insert_sites to generate spreadsheets listing read count, insertion count, and insertion index for each gene. The output files from the tradis_gene_insert_sites analysis were transferred to tradis_comparison.R to compare the reads mapped per gene between the input pools (T0 pools) and the output pools (T12, T24, and T48 pools). Essential genes were determined by analyzing the input library using the tradis_essentiality TraDIS toolkit script (25).

### Construction of isogenic mutant strains

Construction of isogenic mutant strains was performed by previously described methods (31, 34, 35). An internal fragment from six different genes identified as important for fitness in saliva (*spy0644*, *spy0646*, *carB*, *lacR1*, *nifS1*, and *pstS*) was amplified by PCR from genomic DNA from wild-type parental M1 strain MGAS2221 with relevant primers (Table S7), digested with *Bam*HI, and cloned into suicide vector pBBL740 (62). The resulting constructs were transformed into strain MGAS2221 to inactivate each of the six genes. pBBL740 has a *cat* gene which confers chloramphenicol resistance. The plasmid integrant strains (mutants) were selected using THY agar plates supplemented with 5 μg/ml chloramphenicol. The genome of each isogenic strain was sequenced before use to confirm the mutant construct and ensure that no spurious mutations were introduced during mutant construction.

### Saliva growth assay

To compare the ability of GAS strains to grow in human saliva, we first grew GAS strains to mid-exponential phase in THY broth (OD = 0.5). GAS cells were washed three times with PBS, and suspended with the equivalent volume of PBS. 100ul of the GAS-PBS suspension was inoculated into 10 ml aliquots of filter-sterilized human saliva. GAS strains in saliva were incubated at 37° C with 5 % CO_2_. Samples (100 ml) of the GAS-saliva suspension were recovered over time, and GAS CFU was determined by serial dilution and growing on blood agar plates. Four biological replicates were included for each strain. Statistical significance were assessed by repeated-measures 2-way ANOVA.

## ACKNOWLEDGMENTS

This work was supported by funds from the Fondren Foundation (to James M. Musser), the PetPlan Charitable Trust (ref: S14-51) and the Horse Trust (ref: G4104) (to Andrew S. Waller). Amelia R. L. Charbonneau is supported by the University of Cambridge Doctoral Training Partnership scheme, which is funded by the Biotechnology and Biological Sciences Research Council, UK (ref: 1503883). We thank Kathryn Stockbauer for critical reading of the manuscript.

---

**FIG S1** Experimental strategy of TraDIS mutant screens in human saliva.

**TABLE S1** Essential genes identified in the genome of serotype M1 GAS strain MGAS2221 under the conditions tested.

**TABLE S2** Gene mutations conferring decreased fitness after 12, 24 and 48 hour incubation with human saliva.

**TABLE S3** Gene mutations conferring increased fitness after 12, 24 and 48 hour incubation with human saliva.

**TABLE S4** Fitness score of each gene in the M1 GAS genome after 12 hour saliva incubation.

**TABLE S5** Fitness score of each gene in the M1 GAS genome after 24 hour saliva incubation.

**TABLE S6** Fitness score of each gene in the M1 GAS genome after 48 hour saliva incubation.

**TABLE S7** Primers used in this study.

